# Granzyme B is an essential mediator in CD8+ T cell killing of *Theileria parva*-infected cells

**DOI:** 10.1101/325662

**Authors:** Jie Yang, Alan Pemberton, W. Ivan Morrison, Tim Connelley

## Abstract

There is established evidence that cytotoxic CD8+ T cells are important mediators of immunity against the bovine intracellular protozoan parasite *T. parva.* However, the mechanism by which the specific CD8+ T cells kill parasitized cells is not understood. Although the predominant pathway used by human and murine CD8+ T cells to kill pathogen-infected cells is granule exocytosis, involving release of perforin and granzyme B, there is to date a lack of published information on the biological activities of bovine granzyme B. The present study set out to define the functional activities of bovine granzyme B and determine its role in mediating killing of *T. parva*-parasitized cells. DNA constructs encoding functional and non-functional forms of bovine granzyme B were produced and the proteins expressed in Cos-7 cells were used to establish an enzymatic assay to detect and quantify expression of functional granzyme B protein. Using this assay, the levels of killing of different *T. parva*-specific CD8+ T cell clones were found to be significantly correlated with levels of granzyme B protein, but not mRNA transcript, expression. Experiments using inhibitors specific for perforin and granzyme B confirmed that CD8+ T cell killing of parasitized cells is dependent on granule exocytosis and specifically granzyme B. Further studies showed that granzyme B-mediated death of parasitized cells is independent of caspases, but involves activation of the pro-apoptotic molecule Bid.

## Introduction

Antigen-specific CD8+ T cell responses have been shown to play a key role in immunity to a number of viral, bacterial and parasitic infections. One such parasite is the tick-borne protozoan *Theileria parva. T. parva* infects and transforms bovine lymphocytes resulting in an acute, often fatal, lymphoproliferative disease, which is a major constraint to cattle production in a large part of eastern and southern Africa (1). Following invasion of host lymphocytes, the parasite enters the cytosol where it develops to the schizont stage, which triggers a number to signalling pathways that promote host cell proliferation and inhibit apoptosis. By associating with the mitotic spindle of the activated lymphocyte, the parasite is able to divide at the same time as the host cell, ensuring that infection is retained in both daughter cells. Hence, the parasite remains in an intracellular location during this stage of development. Cattle that recover from infection with *T. parva* are solidly immune to subsequent challenge with the same parasite strain but show variable susceptibility to other parasite strains (2). Development of immunity is associated with a potent parasite-specific CD8+ T cell response directed against the parasitized lymphoblasts (3, 4), and transfer of purified CD8+ T cells from immune to naïve twin calves has been shown to confer immunity to parasite challenge (5). The mechanism by which CD8+ T cells mediate protection against *T. parva* is poorly understood. They exhibit strong MHC-restricted cytotoxic activity and secrete IFNγ and TNFα; however, unlike other intracellular protozoa (6, 7), these cytokines do not appear to have a direct effector role against the parasite (8). Hence, cytotoxicity is considered likely to have an important role in immunity, although direct evidence for this is lacking.

As an initial step towards investigating development of subunit vaccines, *T. parva*-specific CD8+ T cell lines have been used successfully to identify a number of target antigens, employing high-throughput screens of expressed parasite cDNAs. Although prime-boost immunisation of cattle with recombinant poxviruses expressing some of these antigens was found to generate specific CD8+ T cell responses, the immunised animals exhibited only partial protection against parasite challenge. A striking feature of the CD8+ T cells induced by this immunisation protocol is that they showed poor cytotoxic activity compared to CD8+ T cells generated by immunisation with live parasites, suggesting poor functional differentiation of the T cell response (9).

Killing of target cells by CD8+ T cells is achieved by release of the contents of secretory lysosomes, known as lytic granules, at the immunological synapse formed upon recognition of class I MHC-bound antigenic peptides by the T cell receptor. Cell killing is initiated by perforin, which creates transient pores in the membrane of the target cell, facilitating uptake into the cytosol of a family of serine proteases known as granzymes. Granzymes exhibit different primary substrate specificities and are able to act on various cellular protein substrates to trigger programmed cell death (10). Five granzymes (A, B, K, H and M) have been identified in humans; mice express four of these granzymes (A, B, K and M) and 6 additional granzymes (C, E, D, F, G and N) (11). We have recently shown that cattle express the same 5 granzymes described in humans, plus a novel granzyme (designated granzyme O) (12). Granzymes have been classified into three distinct evolutionary groups, based on their primary substrate specificities, namely trypsin-like (granzymes A and K), chymotrypsin-like (granzymes B, H, C, E, M, D, F, G and N) and metase-like (granzyme M) (13). The most extensively studied of these proteases, granzyme B, cleaves aspartic acid residues. *In vitro* studies have demonstrated that granzyme B induces target cell death by two main pathways, one involving direct proteolytic activation of caspases (leading to DNA damage) and the other by triggering outer mitochondrial membrane permeabilisation via cleavage of the pro-apoptotic protein, BH3-interaction domain death agonist (Bid) (14). The relative physiological roles of these activities *in vivo* remain unclear, particularly in view of the potential functional redundancy among the granzymes. Nevertheless, gene knockout mice deficient in granzyme B have been shown to have reduced levels of CD8+ T cell-mediated cytotoxicity and have increased susceptibility to some viral infections. Despite the residual ability of CD8+ T cells from granzyme B−/− mice to kill target cells, they were unable to induce DNA fragmentation (15). Extrapolation of findings in mice to other mammalian species is also complicated by the finding of differences in protein substrate specificity between murine and human granzyme B; in contrast to human granzyme B, mouse granzyme B is inefficient at cleaving Bid and is therefore believed to rely largely on direct activation of caspases (16).

In view of the potential importance of CD8+ T cell mediated cytotoxicity as an effector mechanism against *T. parva*, the current study set out to examine the biological activity of bovine granzyme B and to investigate its role in CD8+ T cell-mediated killing of *T. parva*-infected cells. The results demonstrate that granzmye B plays a key role in killing of parasitized cells, that it is able to cleave Bid and that killing occurs predominantly by a caspase-independent pathway.

## Results

### Establishing an *in vitro* assay of granzyme B activity

Bovine granzyme B cDNA incorporated into the pFLAG eukaryotic expression vector (Figure 1A) was expressed in Cos-7 cells and the expressed product tested for enzymatic activity using a substrate assay employing AC-IEPD-pNA, which contains a tetrapeptide recognized specifically by human and murine granzyme B. As shown in Figure 1B, the active form of Granzyme B (pFLAG-Function - with the pro-dipeptide removed) displayed strong activity against the substrate, whereas the native form (pFLAG-WT) and a version containing a mutation in the active tri-peptide site (pFLAG-Mutant) were inactive. As a substrate-specific control, the chymotrypsin substrate Suc-GGF-pNA was used in the assay and no signal was detected with any of the cattle granzyme B constructs.

**Figure 1.**
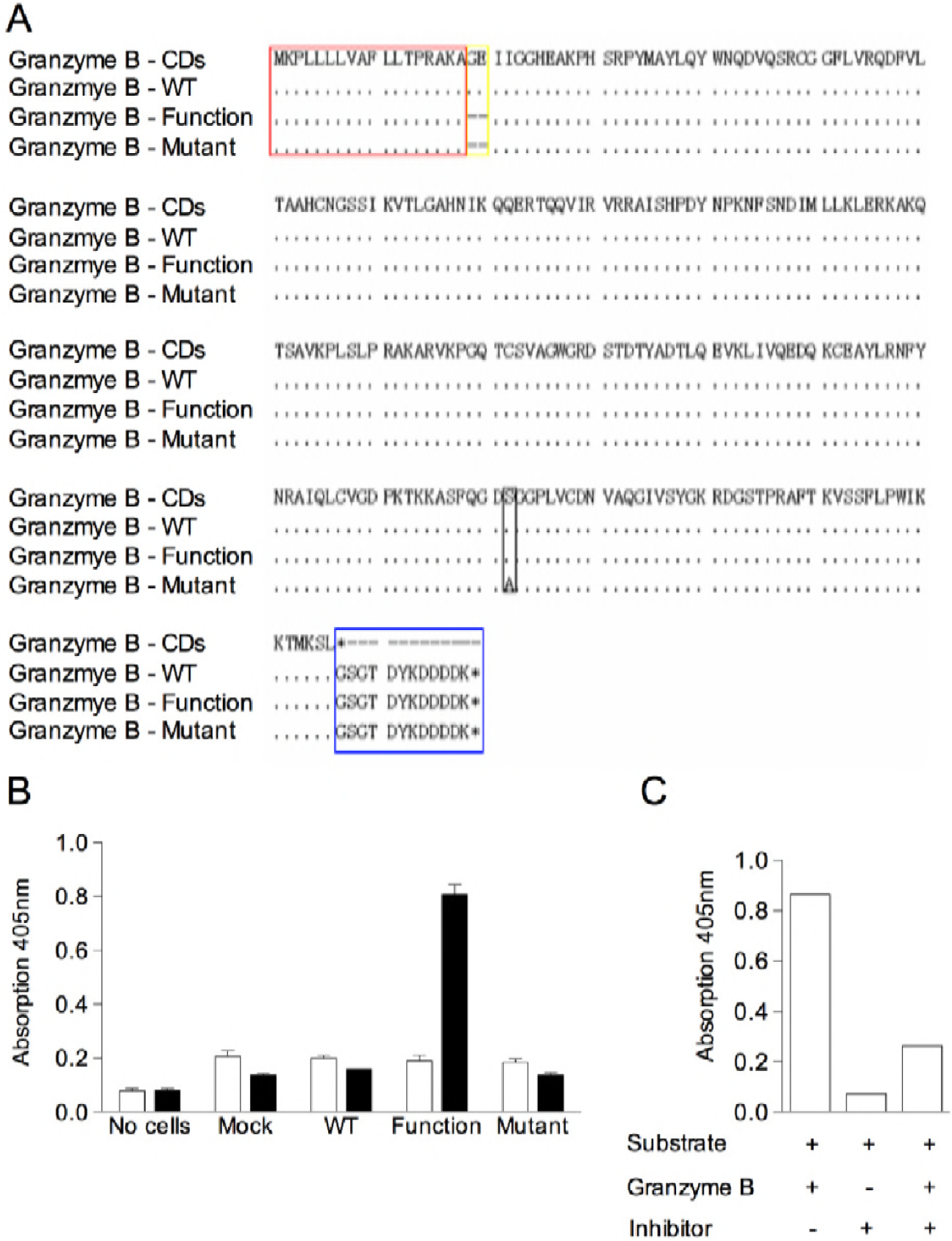
(A). Amino acid sequences from nucleotide sequences of three recombinant forms of bovine granzyme B cDNA, aligned with the reference sequence from the genome database. Granzyme B - CDs - the full length cDNA from bovine genome (corr_ENSBTAG00000010057); Granzyme B - WT - pFLAG-CMVtm-5a vector containing wide type granzyme B; Granzyme B - Function - pFLAG-CMVtm-5a vector containing functional granzyme B; Granzyme B - Mutant - pFLAG-CMVtm-5a vector containing functional granzyme B with Ser_195_ to Ala_195_ mutation. Dot-Identical; Dash-Gap; Leader peptide is highlighted in a red box; Dipeptide/GE is in a yellow box;Ser195Ala is in a black box; FLAG epitope-tag sequence of the pFLAG-CMVtm-5a vector is in blue. (B). Enzymatic activity of different recombinant forms of bovine granzyme B tested on a granzyme B-specific substrate AC-IEPD-pNA (Filled bars) and a control substrate Suc-GGF-pNA (Empty bars): Cos-7 cells were transiently transfected with unmodified granzyme B cDNA (WT), cDNA with the GE dipeptide deleted (Function) or cDNA containing a deletion of the dipeptide and an alanine substitution at position 195 (Mutant). The transfection efficiency of Cos-7 cells with three granzyme B constructs was 35%, 33% and 33%, respectively. Lysates of the transfected cells collected after 48 hours were incubated with the substrates for 4 hours. Controls consisted of lysates of cells transfected with pFLAG without an insert (Mock) and buffer (No cells) added to the substrate. Colour reaction generated after 4 hours by cleavage of the pNA substrate were measured at a wavelength of 405nm using a Synergy^™^ HT Multi-Mode Microplate Reader (BioTek). (C). Inhibition of the functional recombinant cattle granzyme B by preincubating with 10uM granzyme B specific inhibitor AC-IEPD-CHO for 0.5h.

To confirm the specificity of the expressed granzyme B, the enzymatic activity was measured in the presence or absence of the granzyme B inhibitor AC-IEPD-CHO. The specific inhibitor dramatically reduced the activity of the cattle granzyme B preparation by about 4-fold, close to the background level (Figure 1C). The substantial loss of enzymatic function indicates an effective inhibitory capacity of AC-IEPD-CHO for cattle granzyme B.

### Relationship of cytotoxic activity and granzyme B transcript profiles

Analysis of cDNA from *T. parva*-specific CD8+ T cell lines by PCR employing primers that amplify transcripts for 6 defined bovine granzymes demonstrated expression of all six genes, including granzyme B (Figure 2A). The kinetics of granzyme B mRNA expression were examined using a semi-quantitative PCR to determine whether expression was strongly influenced by the time interval after antigenic stimulation. Examination of cDNA prepared from CD8+ T cells at 2-3 day intervals, between 2 and 14 days after stimulation with γ-irradiated *T. parva*-infected cells, demonstrated that near maximal levels of gene expression were achieved between 5 and 7 days after antigenic stimulation, with a subsequent decline in expression (Figure 2B and C). Cells harvested 6-7 days after antigenic stimulation were used for subsequent experiments. To determine whether the levels of killing by CD8+ T cells are related to granzyme B and perforin mRNA expression, CD8+ T cell clones exhibiting different levels of killing were analysed using a semi-quantitative PCR. Two sets of cloned CD8+ T cell lines derived from different animals (641 and 011) were examined; each set of lines expressed identical TCRβ chains and recognised the same epitope but exhibited different levels of cytotoxic activity on autologous parasitized cells (Figure 2D). Of the 4 clones examined from animal 641, two showed no cytotoxic activity whereas the other 2 gave 28-33% killing of parasitized cells. Two of the T cell clones from animal 011 gave high levels of killing (65-75%) of parasitized target cells while the other two clones gave low levels of killing (<30%). Transcripts for granzyme B and perforin were detected in all 8 T cell clones (Figure 2E). Overall, there was no consistent pattern of either granzyme B (r= 0.438, p= 0.278) or perforin (r= −0.104, p= 0.806) mRNA transcript expression that correlated with killing activity (Figure 2F).

### Relationship of cytotoxic activity and level of granzyme B protein expression

A series of CD8+ T cell clones specific for the same epitope in the Tp1 *T. parva* antigen (Tp1_214-224_) were used to examine the relationship between killing activity and granzyme B protein expression. The CD8+ T cell clones showed levels of killing of *T. parva*-infected cells ranging from 1% to 47% at an effector to target ratio of 2:1 (Figure 3A). Granzyme B activity in culture supernatants and in cell lysates of these clones following incubation with infected cells was measured using the *in vitro* substrate-specific assay established above. As shown in Figure 3A, the T cell clones showed variable levels of granzyme B release following exposure to antigen-expressing cells (which prior assays had confirmed do not express granzyme B protein, data not shown). The levels of granzyme activity in cell supernatants showed a close correlation with the levels of granzyme protein in lysates of the respective clones (r= 0.953, p< 0.0001 – Figure 3B), indicating that the levels of enzyme release reflect the cell content rather than inherent differences in rates of release during degranulation. The levels of granzyme B content of the clones showed a statistically significant correlation (r =0.732, p= 0.007) with the levels of cytotoxicity of the T cell clones (Figure 3C).

**Figure 2.**
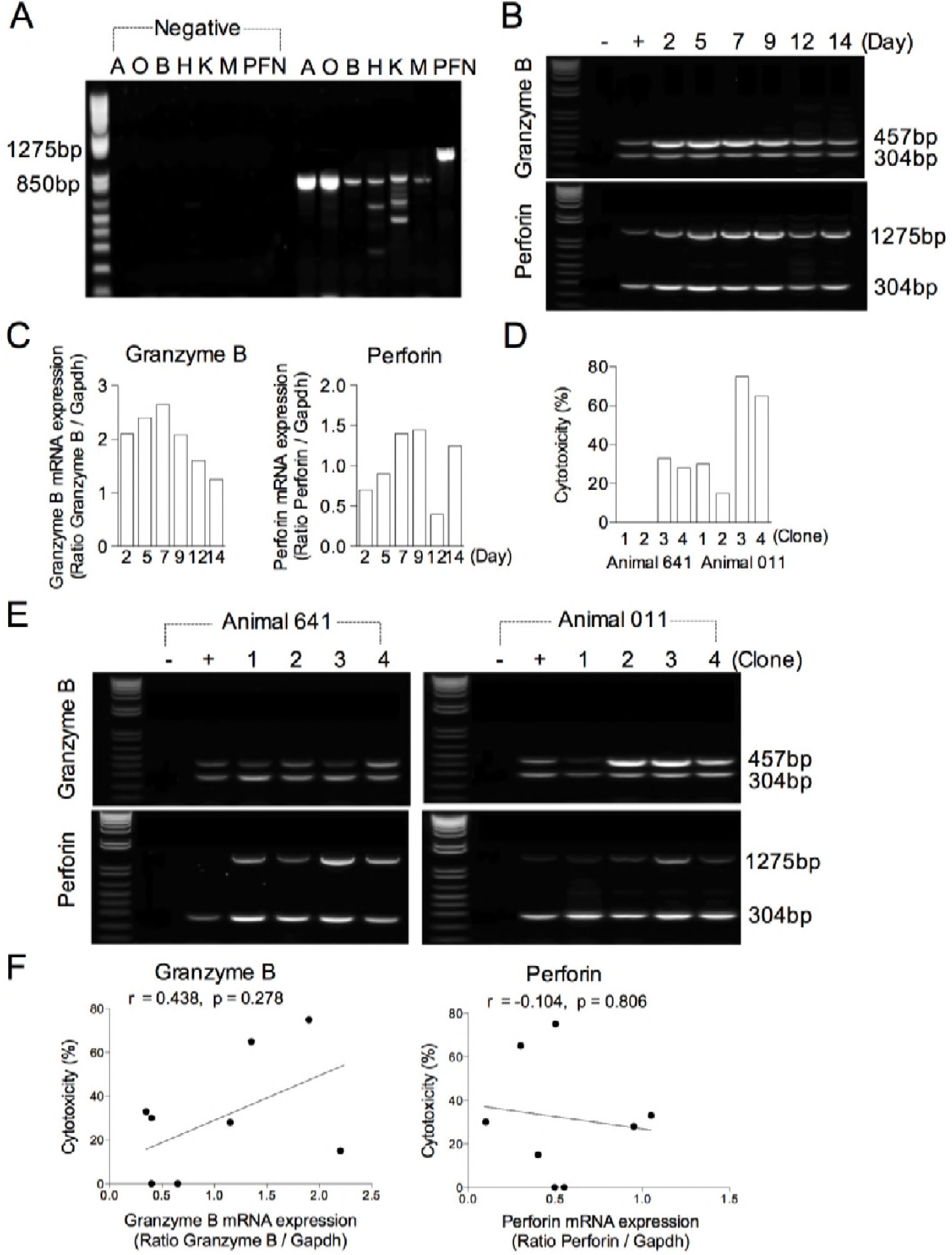
(A). PCR products obtained for each of the bovine granule enzymes from an uncloned *T. parva*-specific CD8+ T cell line (641). The sizes of the PCR products obtained were: granzyme A (A) - 838bp; granzyme O (O) - 849bp; granzyme B (B) - 818bp; granzyme H (H) - 820bp; granzyme K (K) - 889bp; granzyme M (M) - 833bp; Perforin (PFN) - 1275bp; Negative controls (primers with no added cDNA template) were included in the left of the panel. (B). Agarose gels showing the PCR products for granzyme B (457bp), perforin (1275bp) and the GAPDH control (304bp). Days after antigenic stimulation are shown. (C). Changes in quantity of PCR product (vertical axis) at different times following antigenic stimulation, normalised in relation to that of the GAPDH product obtained from the same sample. (D). Cytotoxic activity of 8 *T. parva*-specific CD8+ T cell clones from two different animals (641 and 1011) assayed on autologous *T. parva*-infected targets. (E). Agarose gels showing the PCR products for granzyme B (457bp), perforin (1275bp) and the GAPDH control (304bp) from 8 *T. parva-specific* CD8+ T cell clones (D). (F). Correlation of killing of *Theileria-infected* target cells by CD8+ T cell clones with levels of mRNA expression of granzyme B (r= 0.438, p= 0.278) and perforin (r= −0.104, p= 0.806). Changes in quantity of PCR product (vertical axis) in different T cell clones, normalised in relation to that of the GAPDH product obtained from the same sample. (B, E) A negative control (−), without added template, and a positive control (+), consisting of primers with cDNA template of an uncloned *T. parva*-specific CD8+ T cell line (641) day 7 after 3^rd^ stimulation are included. The density of the all PCR amplicon bands was measured by Kodak 1D software (version 3.6). The correlation between variables was analysed by Pearson’s correlation test. P-values < 0.05 were considered significant.

**Figure 3.**
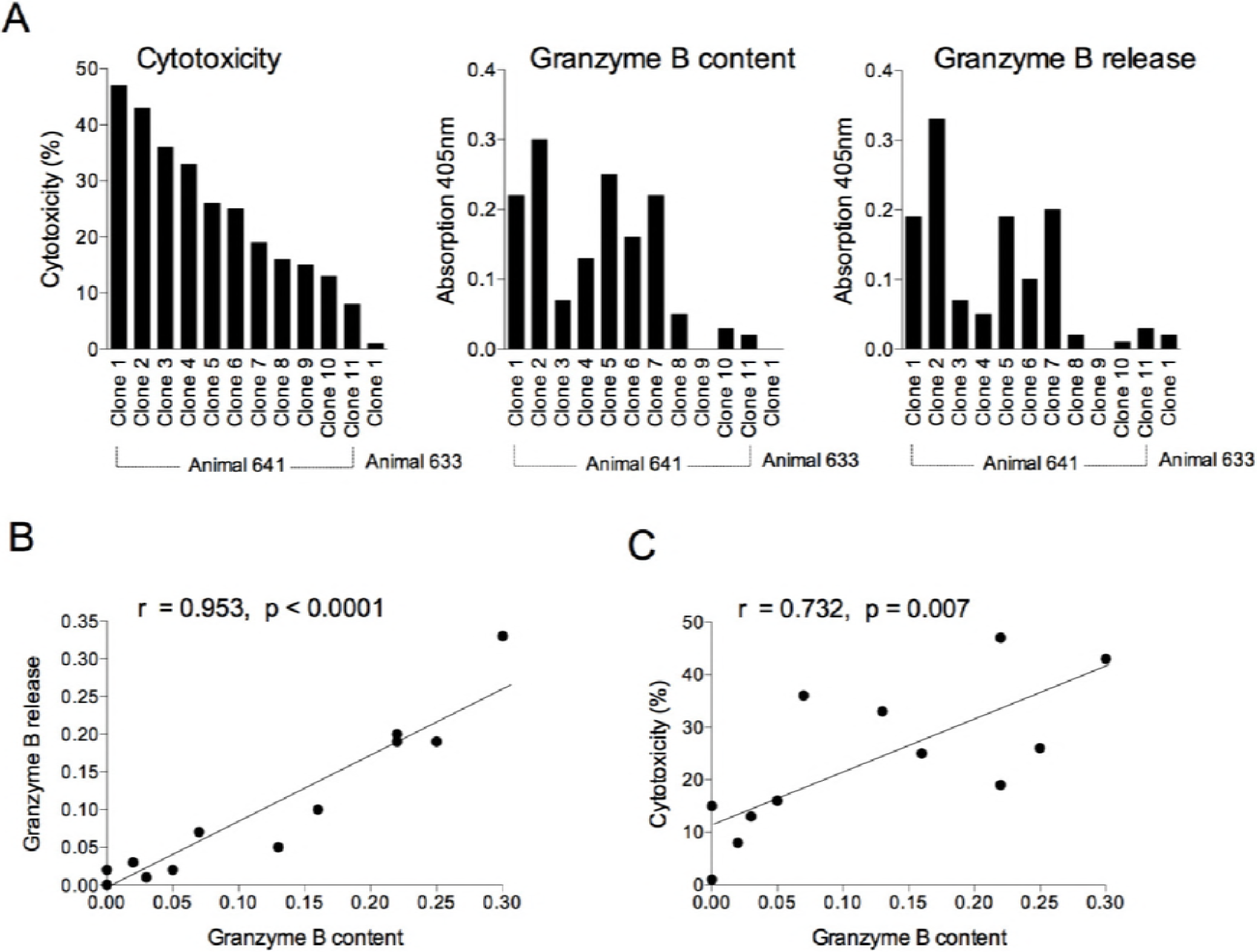
(A). Cytotoxic activity and levels of granzyme B content and release of 12 *T. parva*-specific CD8+ T cell clones isolated from two animals (641 and 633) were assayed with autologous *T. parva*-infected cell target cells. A standard effector to target ratio of 2:1 was used. Correlation of granzyme B cellular activity with (B) levels of released granzyme B following antigenic stimulation (r =0.953, p <0.0001) and, (C) levels of killing of *Theileria-infected* target cells by CD8+ T cell clones (r =0.732, p =0.007). The correlation between variables was analysed by Pearson’s correlation test. P-values < 0.05 were considered significant.

### Cytotoxic activity of T cells is dependent on perforin and granzyme B

The involvement of lytic granule exocytosis and specifically the role of granzyme B in cell killing by bovine CD8+ T cells were investigated by testing the effect of specific inhibitors of perforin and granzyme B. Cytotoxicity assays were first conducted in the presence of a range of concentrations of concanamycin A (CMA), an inhibitor of vacuolar type H^+^-ATPase (17), which raises the pH of the lytic granule and thus induces degradation of perforin (18). The effect of CMA on cytotoxic activity was examined using an un-cloned CD8+ T cell line assayed either on *T. parva*-infected or peptide-pulsed target cells, and 3 cloned CD8+ T cell lines assayed on infected target cells. Concentrations of 10ng/ml or greater of CMA were found to completely ablate killing of all T cell lines (Figure 4A and B). These concentrations of CMA did not affect the viability of the target cells (Figure 4A and B) or the CD8+ T cells (data not shown). The results indicate that lysis of *T. parva*-infected cells by CD8+ T cells is dependent on perforin, implying that killing is mediated by release of granule enzymes.

**Figure 4.**
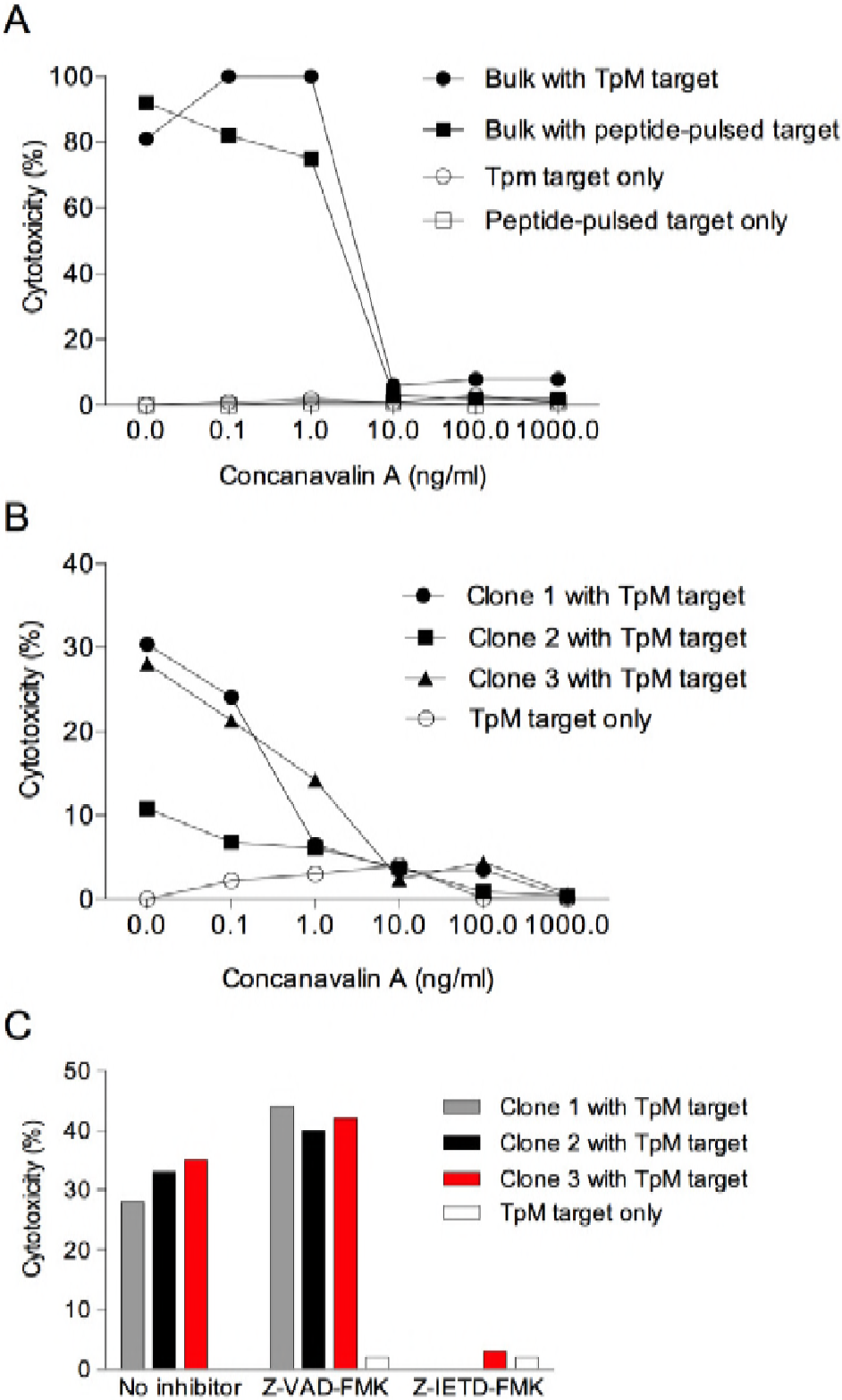
Inhibition of the cytotoxic activity of (A) an un-cloned (‘bulk’) CD8+ T cell line from animal 011 and (B) three CD8+ T cell lines from animal 592 by incubation with the perforin inhibitor concanavalin A (CMA), and (C) three CD8+ T cell lines from animal 641 by incubation with the granzyme B inhibitor Z-IETD-FMK. (A, B) Effectors (1×10^4^) were pre-incubated with various concentrations of CMA for 2h and tested in a 4-h cytotoxicity assay with [^111^In]-labelled autologous TpM target cells and MHC-matched target cells pulsed with Tp2_49-59_ peptide (1000ng/ml). (C) Three cloned CD8+ T cell lines (1×10^4^) were pre-incubated for 1 h with 40uM Z-IETD-FMK and a negative control, Z-VAD-FMK. Labelled target cells alone were also incubated with the inhibitors in the assay. A standard effector to target ratio of 2:1 was used.

To examine the role of granzyme B in cell killing, several specific inhibitors used in studies of murine and human CD8+ T cells were first tested for their ability to inhibit granzyme B activity in bovine CD8+ T cell lysates tested using the *in vitro* substrate-specific assay. Although AC-IEPD-CHO was the most potent inhibitor, reducing granzyme B activity by approximately 80% (Supplementary 1), it is not cell membrane-permeable, prohibiting its use in cellular assays. Z-IETD-FMK, which is membrane permeable and has been shown to inhibit killing of human CD8+ T cells (19, 20), inhibited bovine granzyme B activity by approximately 50% in the substrate assay (Supplementary 1), so was used in subsequent experiments. Pre-incubation of *T. parva*-specific CD8+ T cells with Z-IETD-FMK for one hour prior to use in a cytotoxicity assay resulted in complete inhibition of cytotoxic activity of all 3 cloned T cell lines tested (Figure 4C). A control compound Z-VAD-FMK (a caspase inhibitor that does not affect granzyme B activity of effector cells) did not affect cell killing.

In conclusion, these findings reveal that Z-IETD-FMK specifically and effectively blocks the activity of cattle granzyme B and inhibits killing of target cells by bovine CD8+ T cells - indicating that granzyme B is an important mediator for killing of *T. parva* infected cells.

### Cytotoxic activity of T cells is independent of caspases, but involves activation of Bid

To examine the role of caspases in cell killing, experiments were undertaken to test the ability of the pan-caspase inhibitor Z-VAD-FMK and its control Z-FA-FMK to block killing by two *T. parva*-specific CD8+ T cell clones. In contrast to previous experiments in which this inhibitor was pre-incubated with effector cells (as a negative control), these experiments involved pre-incubation with the target cells. Cytotoxic activity of the CD8+ T cell clones was blocked by inclusion of inhibitors of perforin and granzyme B (CMA and Z-IETD-FMK respectively) but was unaffected by pre-incubation with Z-VAD-FMK (Figure 5A), confirming that killing by these clones was granzyme B-dependent but demonstrating that it was independent of caspase activity. In contrast, Z-VAD-FMK specifically blocked lysis of *Theileria*-infected cells induced by the pro-apoptotic agent cisplatin (Supplementary 2), which is known to mediate cytotoxicity through caspase induction. These results therefore suggest that granzyme B-mediated killing of *Theileria*-infected cells by specific CD8+ T cells is largely independent of caspases.

**Figure 5.**
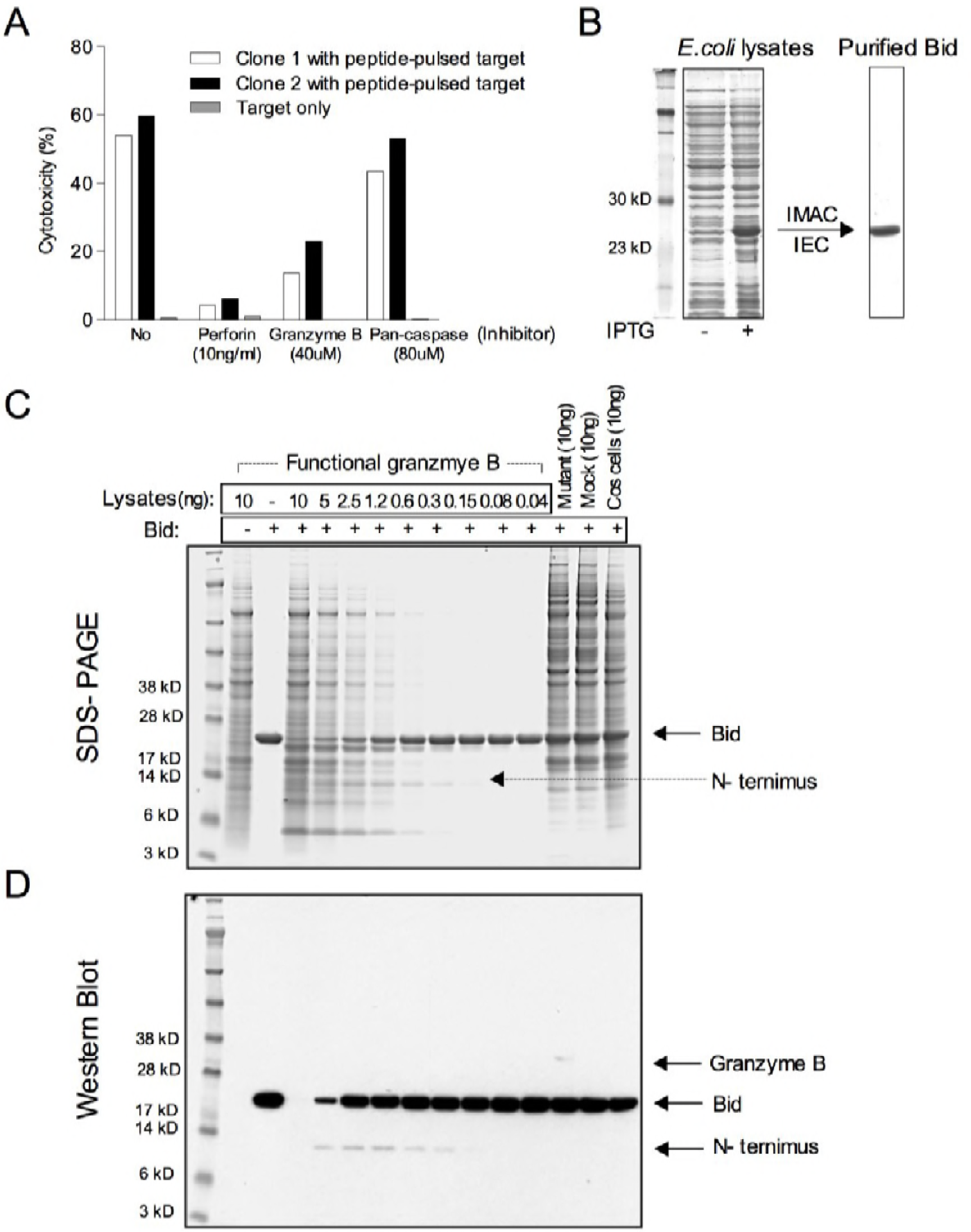
(A). ^111^ In-labelled peptide-pulsed target cells (MHC-matched target cells, 5×10^3^ + Tp1_214-224_, 100ng/ml) were pre-incubated with the ‘pan-caspase’ inhibitor Z-VAD-FMK (80uM) for 1h and tested in a 4-hour cytotoxicity assay with two Tp1-specific cloned CD8+ T cell lines from animal 641. As controls, effector cells (1×10^4^) preincubated with the ‘perforin’ inhibitor CMA (10ng/ml) for 2h or the ‘granzyme B’ inhibitor Z-IETD-FMK (40uM) for 1h were tested in the same experiment. Labelled target cell alone were also incubated with these inhibitors in the assay. A standard effector to target ratio of 2:1 was used. (B). Expression vector pET-15b, carrying an N-terminal His-Tag sequence followed by full-length coding sequence of bovine Bid was expressed in *E. coli* BL21 (DE3) in the presence (+) or absence (−) of IPTG and the expressed products were purified using automated immobilised metal affinity chromatography (IMAC) and automated ion exchange chromatography (IEC). Products were separated by SDS-PAGE and visualized by Commassie blue staining. The predicted size of bovine recombinant Bid is 23.7 kD. (C, D) Purified recombinant bovine Bid proteins (3ug) were incubated with indicated concentrations of active bovine granzyme B for 2 h at 37°C. The reaction products were separated by SDS-PAGE and visualized by Commassie blue staining (C) and full-length recombinant Bid and truncated Bid (N-terminus) were detected by anti-His-Tag antibody and recombinant granzyme B was detected by anti-FLAG M2 antibody in Western Blot (D). (C, D) Inactive bovine granzyme B mutant (an alanine substitution at position 195), mock (pFLAG without an insert) and Cos-7 cells alone were included as negative controls for granzyme B proteolysis specificity

The other mechanism by which granzyme B is known to induce cell death is through cleavage, and so activation, of the pro-apoptotic molecule Bid. To investigate this we sought to examine the ability of bovine granzyme B to cleave bovine Bid. Wild-type bovine Bid was expressed in *E.coli* BL21 with cDNA incorporated into the pET-15b expression vector, which carries an N-terminal His-Tag sequence. Purified recombinant bovine Bid protein (Figure 5B) was incubated with serially titrated concentrations of active bovine granzyme B for 2 hours and the reaction products were separated by SDS-PAGE (Figure 5C). Bovine recombinant Bid was cleaved by active bovine granzyme B as revealed by the detection of an N-terminus fragment of bovine recombinant Bid by an anti-His-Tag antibody in Western Blot (Figure 5D). A reduction in the concentration of bovine granzyme B was associated with a declining ability to cleave bovine recombinant Bid. As controls, an inactive form of bovine granzyme B with a serine to alanine substitution at position 195, mock transfected cells (pFLAG without an insert) and Cos-7 cells alone were analysed; none yielded truncated Bid products, indicating an inability to cleave bovine recombinant Bid. In conclusion, these results indicate that bovine granzyme B mediates cytotoxic effects through the activation of Bid.

## Discussion

In this study we aimed to examine the role of bovine granzyme B in the cytotoxic function of *T. parva*-specific CD8+ T-cell responses. To achieve this we established an *in vitro* substrate-specific assay to detect and quantify expression of bovine granzyme B protein, employing recombinant bovine granzyme B expressed in Cos-7 cells. Using this assay, we showed that the levels of killing of different *T. parva*-specific CD8+ T cell clones are significantly correlated with levels of granzyme B protein and that killing of infected cells by bovine CD8+ T cells is mediated by the granule exocytosis pathway and critically requires granzyme B for induction of cell death. Furthermore, we provided evidence that granzyme B-mediated death of parasitized cells is independent of caspases, suggesting that instead the cell death may be induced via activation of Bid, which we show is cleaved by bovine granzyme B.

Granzyme B was selected for analysis in this study as it has been shown to be the most potent effector molecule utilized by CD8+ T cells to kill infected cells in both humans and mice. Results obtained from generation of recombinant bovine granzyme B and analysis of its biological activity demonstrated many similarities to its human and murine orthologues. This included evidence that processing of the translated polypeptide is similar to that described for humans and mice, with deletion of the dipeptide/G a prerequisite for activation of cattle as well as human and murine granzyme B (21, 22). Similarly, mutation of Ser_195_, one of the functional triad of residues at the conserved catalytic site (His, Asp and Ser), was demonstrated to ablate enzymatic activity of the active form of bovine granzyme B confirming, that as with the murine and human proteins, this residue is a critical component of the enzyme’s active site (23). These similarities extended to the substrate specificities of the human, murine and bovine forms of granzyme B, with recombinant mature bovine granzyme B showing the capacity to cleave AC-IEPD-pNA. This activity forms the basis of a sensitive and reliable *in vitro* method to measure murine and human granzyme B activity (24).

By exploiting this cross-species similarity we were able to generate an equivalent assay for cattle and so investigate levels of biologically active bovine granzyme B and its relation to cytotoxic activity of CD8+ T cells in cattle, overcoming an obstacle posed by the lack of specific antibodies for bovine granzyme B. The demonstration of strong activity against this substrate confirms that cattle granzyme B displays Aspase activity, which is a characteristic feature of granzyme B, with no other known serine protease in mammals having a preference for cleaving Aspartic acid-containing substrates (25). The granzyme B inhibitors AC-IEPD-CHO and Z-IETD-FMK are non-cell-permeable and cell-permeable compounds respectively, which are known inhibitors of human and rodent granzyme B (19, 26). Herein, we have demonstrated that these compounds also effectively inhibit bovine granzyme B, further highlighting the cross-species functional similarities. However, the inability of another two inhibitors of human and murine granzyme B (Z-AAD-CMK and AC-AAVALLPAVLLALLAPIETD-CHO) to block bovine granzyme B (data now shown) emphasises that extrapolating functional parameters based on orthology can’t be assumed for granzymes and must be empirically validated.

This also applies to the pathways utilised by granzyme B to mediate killing, which are known to be species-dependent. Mouse granzyme B predominantly functions through the direct activation of caspases to promote apoptosis, whereas human granzyme B acts mainly via a Bid-dependent pathway (16, 27). Consequently, the mechanisms used by bovine granzyme B to induce cell death could not be implied based on cross-species extrapolation. Work described in this study demonstrates that bovine granzyme B, like its human orthologue, is capable of cleaving Bid protein *in vitro*, thus providing evidence indicating that Bid activation can be utilised by bovine granzmye B for cell death induction. Although activation of caspases was initially thought to be important in granzyme B-mediated cell death, studies by many groups revealed that requirement for caspase activation, even in mice, isn’t absolute. For example, an *in vitro* study of mouse CD8+ T cells showed that apoptotic nuclear damage induced by granule exocytosis was abrogated by the caspase inhibitor Z-VAD-FMK, whereas lysis of the cells was unaffected. In contrast, target cell lysis induced by the pro-apoptotic drug cisplatin was specifically blocked by this inhibitor (28). Similar results have been obtained in studies with purified human granzyme B; caspase inhibition preventing granzyme-induced DNA damage but not cell lysis (29). These observations are consistent with the results obtained in this study, which showed that Z-VAD-FMK inhibited cisplatin-induced apoptosis of *Theileria*-infected cells, but did not inhibit granzyme B-mediated cytolytic activity of cattle CD8+ T cells. *T. parva* has been shown to protect infected cells from apoptosis by utilizing *NF-kB* activation to induce the expression of anti-apoptotic proteins such as FLIP (which functions as a catalytically inactive form of caspase-8), X-chromosome-linked inhibitor of apoptosis protein (XIAP) and c-IAP (which block caspase-9 and also downstream executioner caspases 3 and 7) (30). Studies by Guergnon in 2003 showed that drug-induced parasite death in *Theileria*-infected cells resulted in apoptosis involving activation of caspases 9 and 3 and was inhibited by Z-VAD-FMK (31). These findings confirmed that bovine caspases in non-granzyme B mediated killing are capable of inducing cell death and that Z-VAD-FMK is an effective inhibitor of bovine caspases. The inhibition of killing by *T. parva*-specific CD8+ T cell clones by Z-IETD-FMK but not Z-VAD-FMK in the current study demonstrates that T cell-mediated killing of *T. parva*-infected cells is dependent on granzyme B but independent of caspases. Although this may be universally applicable to bovine granzyme B mediated cytotoxicity, it is important to note that as a consequence of the negative regulation of caspases by intracellular inhibitors induced by the *NF-κB* pathway in *T. parva*-infected cells the apparent redundancy of caspases might be a feature of this specific biological context.

The prime rationale for conducting this study was to better understand the molecular mechanisms that underlie the functional capacity of *T. parva*-specific CD8+ T-cells. The critical role that these cells play in mediating immunological protection against *T. parva* (29) has led to considerable efforts to identify CD8 T cell target antigens for use in generating novel subunit vaccines (32, 33). A number of *T. parva* antigens recognised by CD8 T cells from immune cattle have been identified and, although they have proved to be immunogenic when used in prime-boost immunisation protocols, the CD8+ T-cells elicited have generally exhibited poor cytotoxicity and have been demonstrated to be poorly protective upon *in vivo* challenge (9). Understanding the discrepancy between immunogenicity and protective efficacy will be critical to defining the ‘correlates of protection’ that can guide subsequent vaccine development. Ongoing work is applying transcriptomics to address this issue, however such approaches used in isolation have limitations.

Our data, from assays of expressed biologically active granzyme B, revealed a statistically significant correlation between the levels of granzyme B enzymatic activity in cell lysates (and supernatants) of cloned CD8+ T cell lines and levels and killing of *T. parva*-infected cells, indicating that granzyme B is a dominant effector molecule in CD8+ T-cell mediated killing of these parasitized cells. A prominent role for granzyme B was corroborated by subsequent analysis showing that the membrane-permeable inhibitor of granzyme B, Z-IETD-FMK, reduced *T. parva*-infected cell lysis by these CD8+ T-cells by 70-100%. However, the correlation was not absolute. For example, one clone consistently showed low levels of granzyme B content and release but displayed relatively strong killing (Figure 3A). Such discrepancies suggest that additional factors, such as other granzymes, that may vary between T cells clones also contribute to the cytotoxic activity of these CD8+ T cells. There is substantial evidence from studies in humans and mice that other granzymes can effectively mediate cell death by themselves and/or synergistically increase the activity of granzyme B *in vitro.* Examples from the literature include i) co-transfection of rat basophilic leukemia (RBL) cells with granzyme A and granzyme B in the presence of perforin resulting in enhanced killing of tumour targets in a synergistic manner (34); ii) human granzyme H can augment granzyme B-mediated killing of adenovirus-infected cells (35–37) by neutralizing the viral inhibitor of granzyme B (L4-100K assembly protein) (36, 37) and iii) human granzyme M can rapidly induce cell death of tumor cells *in vitro* directly (38–40) as well as hydrolyse PI-9, thereby inactivating its inhibitory function for granzyme B (41). Such examples illustrate the potential for other bovine granzymes to cooperate with granzyme B in achieving CD8+ T cell-mediating cell death of *T. parva*-infected cells. Unfortunately, further investigation of these interactions in cattle is hampered by the current lack of specific antibodies and biological assays to measure other bovine granzyme proteins. However, this study, which provides the first description of the biological activity of a member of the granzyme family in cattle, exemplifies an approach that could readily be applied to study other bovine granzymes.

In conclusion, work described in this paper has developed molecular and biochemical methods to define the functional activities of bovine granzyme B and demonstrated an indispensible role for granzyme B in killing of *T. parva*-infected cells by specific CD8+ T cells. This study represents the first dissection of the effector mechanisms employed in killing of target cells by bovine CD8+ T cells and specifically provides the first evidence that granzyme B plays a key role in killing of *T. parva*-infected cells by specific CD8+ T cells.

## Materials and Methods

### Animals and T cell lines

Four Holstein-Friesian animals (011, 592, 641 and 633) homozygous for the A10 or A18 MHC I haplotypes were used for the study. Their MHC types were determined by a combination of serological typing (42) and MHC I allele-specific PCR (43). The animals were aged 18-36 month at the outset of the study and were maintained indoors on rations of hay and concentrate. Cattle were immunized against the Muguga stock of *T. parva* (TpM) by infection with cryopreserved sporozoites and simultaneous administration of a long-acting formulation of oxytetracycline as described previously (2). Animals were challenged with a lethal dose of sporozoites on two occasions at ~18-month intervals following immunization. All animal experimental work was completed in accordance with the Animal (Scientific Procedures) Act 1986. *T. parva*-specific CD8+ T cell lines and clones were generated from the immune cattle and maintained as described previously (44).

### Standard and semi-quantitative PCR assays

Total RNA was extracted from *T. parva*-specific CD8+ T cell lines from immunized cattle using Tri-reagent (Sigma) and cDNA was synthesised using the Reverse Transcription System (Promega) with priming by the Oligo (dT)15 primer, both according to the manufacturer’s instructions. The primers for granzymes and perforin and the protocols for standard PCR reactions were as previously described (12). For semi-quantitative PCR, the sequences of primers were as follows: granzyme B: 5’-ACT GGA ATC AGG ATG TCC AGA G-3’ (Forward), 5’-TTT GGG TCC CCC ACA CAC AG-3’ (Reverse) and Gapdh: 5’-ACC CCT TCA TTG ACC TTC AC-3’ (Forward); 5’-TTC ACG CCC ATC ACA AAC ATG-3’ (Reverse). The PCR reactions were composed of 20pmol of granzyme B/perforin primers and 10pmol of Gapdh primers, 2.5 units BIOTAQ (5 units/ul, Bioline), 2.5ul SM-0005 buffer, 0.05ug of cDNA template and nuclease-free water to give a final volume of 25ul. The primers for perforin and the protocol for PCR programme were as described above. Semi-quantified PCR products were analysed by 1.5% agarose gel electrophoresis and the density of the specific bands was measured by computer software (KODAK 1D 3.6 version).

### Cloning of bovine granzyme B cDNA constructs

Full-length bovine wild-type (WT) granzyme B was amplified from cDNA by high fidelity PCR, using primers flanking the coding sequence as previously described (12). The high fidelity PCR protocol was composed of 10pmol of primers, 1.2 unit *Pfu* DNA polymerase (3 units/ul, Promega), 10x Buffer with MgSO_4_ (Promega), 10mM dNTP, 0.5ug of cDNA template and nuclease-free water to give a final volume of 50ul. The programme used was as follows: 95°C for 2 min, 30 cycles of 95°C for 1min followed by 55°C for 0. 5 min and 72°C for 2.5 min, and a final extension period of 72°C for 5 min. To generate cDNA encoding active granzyme B, six nucleotides encoding a dipeptide segment in the wild-type granzyme B cDNA (which inhibits granzyme B function and is present in pro-granzyme B but absent in fully mature granzyme B) were deleted by PCR splice overlap extension (PCR-SOE), based on procedures described for human granzyme B (21). Briefly, two PCR assays were initially performed to generate two overlapping fragments that carry the six-nucleotide deletion in the overlapping segment. These reactions utilised the external flanking primers described above with the following internal primers: 5’-CAAAGGCAATCATCGGGGGCCATG-3’ (Forward); 5’-CCCGATGATTGCCTTTGCCCTGGG-3’ (Reverse). The resulting two fragments were mixed, denatured and annealed to produce deletion mutant DNA templates and amplification of the extended DNAs was performed with flanking primers in a further PCR. Substitution of Ser with Ala at the active site of dipeptide-knockout cDNA was performed by ‘megaprimer’ PCR mutagenesis (45, 46) using an internal mutagenic forward primer incorporating the mutation as follows: 5’-AGAAAGCTTCCTTTCAGG GGGAC<u>G</u>CGG-3’. Briefly, an initial 5 cycles of a PCR reaction containing 50pmol of internal mutagenic forward primer and 2.5pmol of a flanking reverse primer (as described above) was followed by a prolonged extension step to generate mutant mega fragments. 50pmol of the other flanking primer (as described above) was added to the mutant templates and the PCR reaction subjected to a further 25 cycles to generate full-length product containing the mutation. All three bovine granzmye B cDNAs were sub-cloned into the pFLAG-CMV^tm^-5a expression vector (Sigma) and nucleotide sequencing performed by DBS Genomic (Durham University).

### Expression of granzyme B in Cos-7 cells

Cos-7 cells were maintained in Dulbecco’s Minimal Essential Medium (DMEM, Invitrogen) supplemented with 10% FCS, 5×10^−5^ M 2-Mercaptoethanol, 4mM glutamine, 100U/ml penicillin and 100ug/ml streptomycin, at 37°C with 5% CO_2_. Cos-7 cells were transfected in 75cm^2^ flasks with the pFLAG-CMV^tm^-5a vector containing each of the three cattle granzyme B recombinant cDNAs (wild type (60ug), dipeptide knockout (60ug) and knockout with a Ser195Ala substitution (40ug)) or vector only (60ug). The transient transfection was performed by using the Lipofectamine^tm^ 2000 reagent (Invitrogen) according to the manufacturer’s protocol. Transfected cells were harvested after 48h, washed and suspended in cold PBS and analysed in further experiments. To test for expression of transfected DNA products, cytospin smears of cells were examined microscopically with anti-FLAG M2 antibody (1:500 dilution; IgG1; Sigma). The transfection efficiency of pFLAG vectors containing three bovine granzyme B recombinant cDNAs containing WT, the dipeptide knockout and the knockout with an additional Ser195Ala substitution was 35%, 33% and 33%, respectively.

### Granzyme B protease activity in transfected Cos-7 cells

Cell lysis and assay of protease activity were performed as previously described for equine granzyme B (47). Briefly, aliquots of 1ml of PBS-washed Cos-7 cells adjusted to 2×10^6^ cells/ml in PBS were pelleted and lysed by addition of 0.2ml lysis buffer (1%Triton X-100, 50mM Tris, pH8.0 and 2ul of Benzonase Nuclease 25U/ml, Purity>99%, Merck). Following incubation on ice for 20min, lysed cells were centrifuged at 21,000 × g for 10min at 0°C to pellet cell nuclei and other cell debris. Supernatants were harvested and assayed in duplicate for protease activity; aliquots of 25ul of lysis supernatant, granzyme B substrate Ac-IEPD-pNA, (Calbiochem) at a final concentration of 300uM and reaction buffer (0.1M Hepes, pH 7.0; 0.3M NaCl; 1mM EDTA) in a total volume of 250ul/well were added into the wells of Falcon™ 96-Well Flat bottomed Microplates (BD). The chymotrypsin substrate I, Suc-GGF-*p*NA (Calbiochem), was used as a negative control for substrate specificity. The reaction was composed of 25ul of lysis supernatant, 1mM Suc-GGF-pNA in the final concentration and the reaction buffer (50mM Tris, 100mM NaCl, pH8.0) in a total volume of 125ul. Mixtures were incubated at 37°C for 4h and the colour reaction generated by cleavage of the pNA substrate measured at a wavelength of 405nm by using a Synergy™ HT MultiMode Microplate Reader (BioTek). For inhibition of active bovine granzyme B protease activity in lysates, aliquots of 25ul of lysis supernatant containing active bovine granzyme B were pre-incubated with 10uM Ac-IEPD-CHO (the granzyme B inhibitor, Calbiochem) at 37°C for 0.5h.

### Granzyme B activity in CD8+ T cell lines

Methods used for measurement of granzyme B in T cell lysates and supernatants were based on procedures previously described for human and equine granzyme B (24, 47). CD8+ T cells washed in PBS were adjusted to 1×10^6^ cells/ml in PBS, pelleted and lysed by addition of 50ul of a lysis buffer per ml as described above. To examine granzyme B release, aliquots of 1×10^6^ CD8+ T cells were distributed into the wells of 96-well V-bottomed plates together with 5×10^5^ target cells in a total volume of 200ul phenol-red-free complete media (RPMI 1640 with 5% FCS, Invitrogen,). Control wells containing effector cells and medium were also included. After incubation in an atmosphere of 5% CO_2_ at 37°C for 4h, the plates were centrifuging for 10min at 400xg and supernatants were collected. Granzyme B activity was measured by adding aliquots of 10ul of cell lysates or 40ul of culture supernatants in duplicate to wells of Falcon™ 96-well flat-bottomed Microplates (BD) together with 200uM granzyme B substrate, Ac-IEPD-pNA, (Calbiochem) and reaction buffer (0.1M HEPES, pH7.0; 0.3M NaCl; 1mM EDTA) in a total volume of 100ul/well. Wells containing reaction buffer and substrate control were also included as controls. Mixtures were incubated at 37°C for 4h and the colour reaction generated by cleavage of the pNA (p-nitroaniline) substrate measured at a wavelength of 405nm using a Synergy™ HT Multi-Mode Microplate Reader (BioTek). To test for specificity of the reaction, CD8+ T cells were preincubated with the cell-permeable granzyme B inhibitor, Z-IETD-FMK (40uM) for 1h prior to preparation and testing of cell lysates as describe above. 40uM Z-VAD-FMK, a pan-caspase inhibitor was used as a negative control, whereas a non-cell-permeable granzyme B inhibitor, AC-IEPD-CHO (10uM) was used to inhibit granzyme B activity in lysates as a positive control.

### Cytotoxicity assays

Standard 4-hour [^111^In]-release cytotoxicity assays were used to measure cytotoxicity of CD8+ T cell clones, using as target cells either autologous *T. parva*-infected cells or autologous *T. annulata* - transformed cells incubated with peptide for 0.5h prior to the assay (44). Peptides were supplied by Pepscan Systems (Lelystad, The Netherlands). All assays were conducted in duplicate, and controls included *T. annulata*-infected target cells without added peptide and, where appropriate, MHC-mismatched *T. parva*-infected target cells. Cytotoxicity assays were established and specific lysis was measured as described previously (44). For inhibition of perforin activity, effector cells were pre-incubated with ten-fold dilutions of concanamycin A (CMA) at final concentrations ranging from 0.1ug/ml to 1000ug/ml for 2h at 37 °C. For inhibition of granzyme B activity, effector cells were pre-incubated for 1h at 37°C with 40uM Z-IETD-FMK and the negative control, pan-caspase inhibitor Z-VAD-FMK (40uM). For inhibition of caspase activity, ^111^In labelled target cells were pre-incubated with 80uM Z-VAD-FMK and the negative control, cathepsin B Inhibitor Z-FA-FMK (80uM) for 1h at 37°C.

### Generation of recombinant bovine Bid

Wild-type bovine Bid cDNA was amplified using primers flanking the full-length coding region of bovine Bid as follows: 5’-GAAGCTTAGCATATGGATTTGAAGGTTAGC- 3’ (Forward); 5’-TTCTGCCGAGGATCCACTCAGTCCATCTGATTTCGG- 3’ (Reverse). The amplified PCR products were purified and sub-cloned into the NdeI and BamHI sides of pET-15b vector (Novagen) and nucleotide sequencing performed by DBS Genomic (Durham University). The protocols for expression and purification of recombinant bovine Bid proteins were performed as previously described for human Bid (48). Briefly, pET-15b expression vectors containing wild-type bovine Bid cDNA were transformed in *E.coli* BL21 (DE3) pLYsS (Novagen) and expressed in the presence of IPTG. The expressed products, which carry an N-terminal His-Tag sequence, were purified with automated immobilised metal affinity chromatography (IMAC) using a nickel affinity column (Qiagen) and further purified with automated ion exchange chromatography (IEC) using a Mono Q column (Pharmacia).

### Proteolysis of recombinant bovine Bid by bovine granzyme B

Two-fold dilutions of lysates containing active bovine granzyme B at final concentrations ranging from 10ng to 0.04ng in 10ul reaction volumes were incubated with 3ug of recombinant bovine Bid for 2h at 37°C. Inactive mutated bovine granzyme B (an alanine substitution at position 195), mock (pFLAG without an insert) and Cos-7 cells alone were used as negative controls for granzyme B proteolysis specificity. Reaction products were separated by SDS-PAGE (NuPAGE 4-12% Bis-Tris gel, Thermo Fisher) and visualized by Coomassie blue staining. The reaction products were transferred using the iBlot (Thermo Fisher) for Western blotting, according to the manufacturer’s instructions. The blots were probed with anti-His Tag antibody (1:2500 dilution, Thermo Fisher) and anti-FLAG M2 antibody (1:1000 dilution, Sigma) and detected by chemiluminescence using HRP-labelled rabbit anti-mouse IgG (H+L) secondary antibody (1:5000 dilution, Thermo Fisher).

### Statistical analysis

Statistical analyses were performed using Minitab software (Minitab® 15.1.20.0, Minitab Inc.). The correlation between variables was analysed by Pearson’s correlation test. P-values < 0.05 were considered significant.

## Acknowledgements

This work was supported in part by a grant awarded by the Bill and Melinda Gates Foundation jointly with the UK Department for International development (DfID) [No. OPP1078791] and a Biotechnology and Biological Sciences Research Council (BBSRC) Institute ISP grant (grant number BB/J004227/1). Jie Yang was supported by a China Scholarship Council/University of Edinburgh Scholarship. The authors also wish to thank Kathryn Degnan and Robyn Cartwright for expert technical assistance, Dr Andy Gill for technical assistance with protein purification and analysis and Dr Darren Shaw for statistical help and advice.

---

**Supplementary 1.**
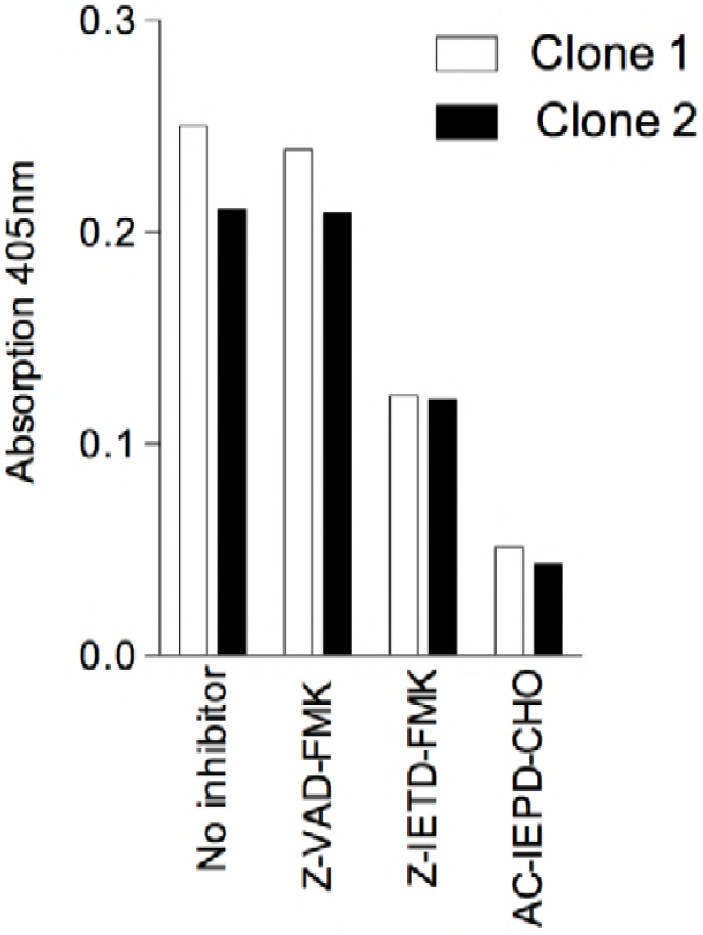
. Effect of granzyme B inhibitors on granzyme B enzymatic activity in *T. parva*-specific CD8+ T cell lines. Two cloned CD8+ T cell lines (1×10^6^ cells) were pre-incubated for 1h with the cell-permeable granzyme B inhibitor, Z-IETD-FMK (40uM) and a negative control, Z-VAD-FMK (40uM) and tested in a 4-hour substrate assay. As a positive control, lysates of CD8+ T cell lines (1×10^6^ cells) were also pre-incubated with the non-cell-permeable granzyme B inhibitor, AC-IEPD-CHO (10uM) and tested in the same substrate assay.

**Supplementary 2.**
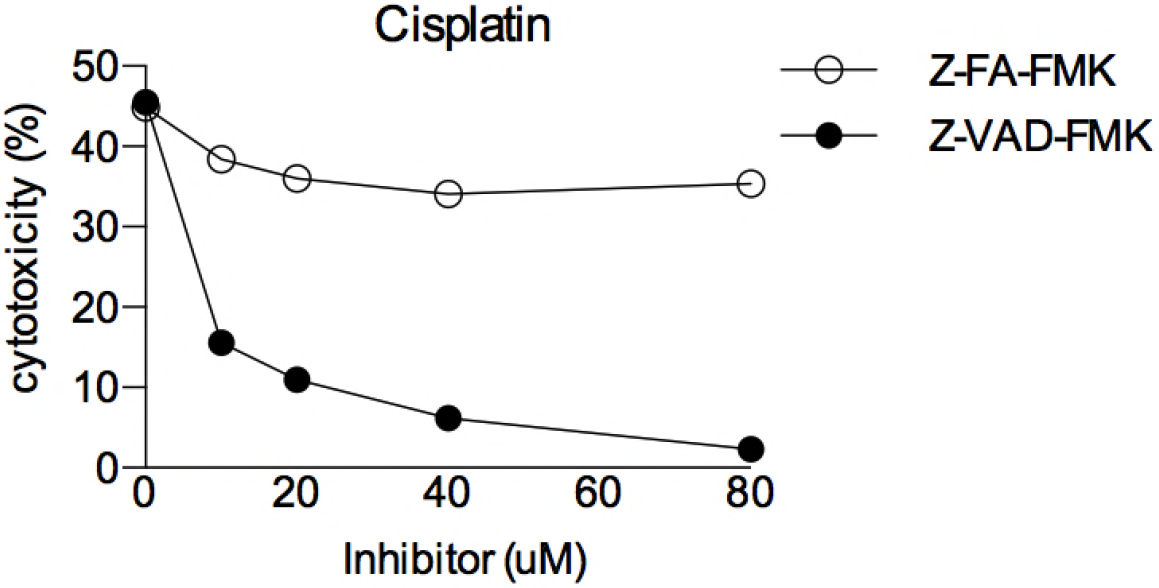
Inhibition of cytolysis of *T.annulata*-infected cells induced by 100uM cisplatin by incubation with Z-VAD-FMK for 24 hour. ^111^ In-labelled *T. annulata*-infected cells (5×10^5^) from animal 641 were incubated for 24h with 100uM cisplatin with various concentrations of Z-VAD-FMK and radioactivity released was measured. A negative control inhibitor, Z-FA-FMK was included. Labelled target cells incubated with Z-VAD-FMK (80uM) were used to measure spontaneous radioactivity released.

